# Top-down control of the descending pain modulatory system drives placebo analgesia

**DOI:** 10.1101/2025.02.13.638185

**Authors:** Giulia Livrizzi, Jingzhu Liao, Desiree A. Johnson, Susan T. Lubejko, Janie Chang-Weinberg, Chunyang Dong, Lin Tian, Matthew R. Banghart

## Abstract

In placebo analgesia, prior experience and expectations lead to pain suppression by the administration of an inert substance, but causal evidence for its neural basis is lacking. To identify the underlying neural circuits, we reverse-translated a conditioned placebo protocol from humans to mice. Surprisingly, the placebo effect suppresses both nociception and unconditioned emotional-motivational pain-related behavior. Descending pain modulatory neurons in the periaqueductal gray (PAG) are critical for both morphine and placebo antinociception. The placebo effect depends on input to the PAG from the medial prefrontal and anterior cingulate cortices, but not anterior insular cortex. Conditioning enhances noxious stimulus-evoked endogenous opioid release in the PAG to produce analgesia. Our results suggest that cortical control of the descending pain modulatory system (DPMS) is gated by rapid endogenous opioid signaling in the PAG during placebo trials. This study bridges clinical and preclinical research, establishing a central role for the DPMS in placebo analgesia.

## Main

Placebo analgesia is a form of cognitive pain modulation that can result from behavioral conditioning and expectations about noxious stimuli^1–5^. Placebo effects contribute to the efficacy of analgesic drugs but can obscure the effects of therapeutics in clinical trials^5,6^. Understanding the neurobiology of placebo analgesia might enable the underlying mechanisms to be harnessed for clinical therapies, potentially reducing reliance on opioid drugs^7,8^. Functional imaging studies in humans have revealed activity changes in prefrontal, anterior cingulate, and insular cortices during placebo analgesia, as well as subcortical structures such as the periaqueductal gray^9–12^. Endogenous opioid neuropeptide signaling appears to be critical, as most forms of placebo analgesia are blocked by opioid antagonists^4,13,14^. Furthermore, positron emission tomography studies with radiolabeled opioid drugs have identified putative sites of peptide release^15,16^.

Strikingly, many of the brain regions implicated in placebo analgesia are also activated by exogenous opioid analgesics such as morphine, suggesting that placebo effects occur through a conserved pain modulatory network^17^. However, because our current understanding of placebo circuits is primarily based on correlative measures of neural activity and neurochemical signaling in human subjects, it has been challenging to ascribe causal roles to the underlying neural pathways^9,10,18,19^. Furthermore, several studies have drawn into question many of conclusions based on early, underpowered analyses, in particular the notion that placebo analgesia involves suppression of nociception through the descending pain modulatory system (DPMS)^20–23^. The neural circuit mechanisms of placebo analgesia are important to resolve if therapeutic strategies are to be developed that capitalize on placebo-like processes to reduce pain.

### Morphine-conditioned placebo analgesia in mice

In order to study the circuit mechanisms of placebo analgesia in mice, we established reliable placebo antinociception protocols based on contextual conditioning with morphine, under the premise that contextual cues associated with morphine pain relief might promote pain suppression in the absence of drug treatment. Morphine conditioning has proven highly effective at producing placebo pain relief in human subjects^4,24^. Although not studied intensively, it has also been shown to produce antinociception in rodents^14,25–27^. In our placebo protocols, mice were injected with saline or morphine, and then placed into contexts distinguished by both visual and olfactory cues (**Figure 1A**). After a 30 min wait period to achieve maximal morphine antinociception, paw withdrawal latencies were measured on the hot plate as a metric of thermal pain sensitivity. Importantly, mice were removed promptly after exhibiting the first paw withdrawal to prevent sensitization to the assay. After conditioning, the placebo trial on test day consisted of saline administration and placement in the morphine-paired context prior to the hot plate test. In protocol 1, male mice were conditioned over four days with two exposures each to either morphine (10 mg/kg, *i.p.*) or saline (**Figure 1B**). This conditioning protocol produced strong placebo antinociception (52.4± 10.0%), quantified as the percent of the paw withdrawal latency increase produced by the prior dose of morphine (**Figure 1C,D**; average of 3 consecutive trials is shown for each subject). In a subset of mice, subsequent administration of saline in the saline context showed that the placebo effect is context-specific. We also categorized subjects as either responders or as non-responders (see methods) (**Figure 1E**). Considering responders only, this placebo protocol produced 60.5±6.8% of the morphine-induced antinociception.

**Figure 1:**
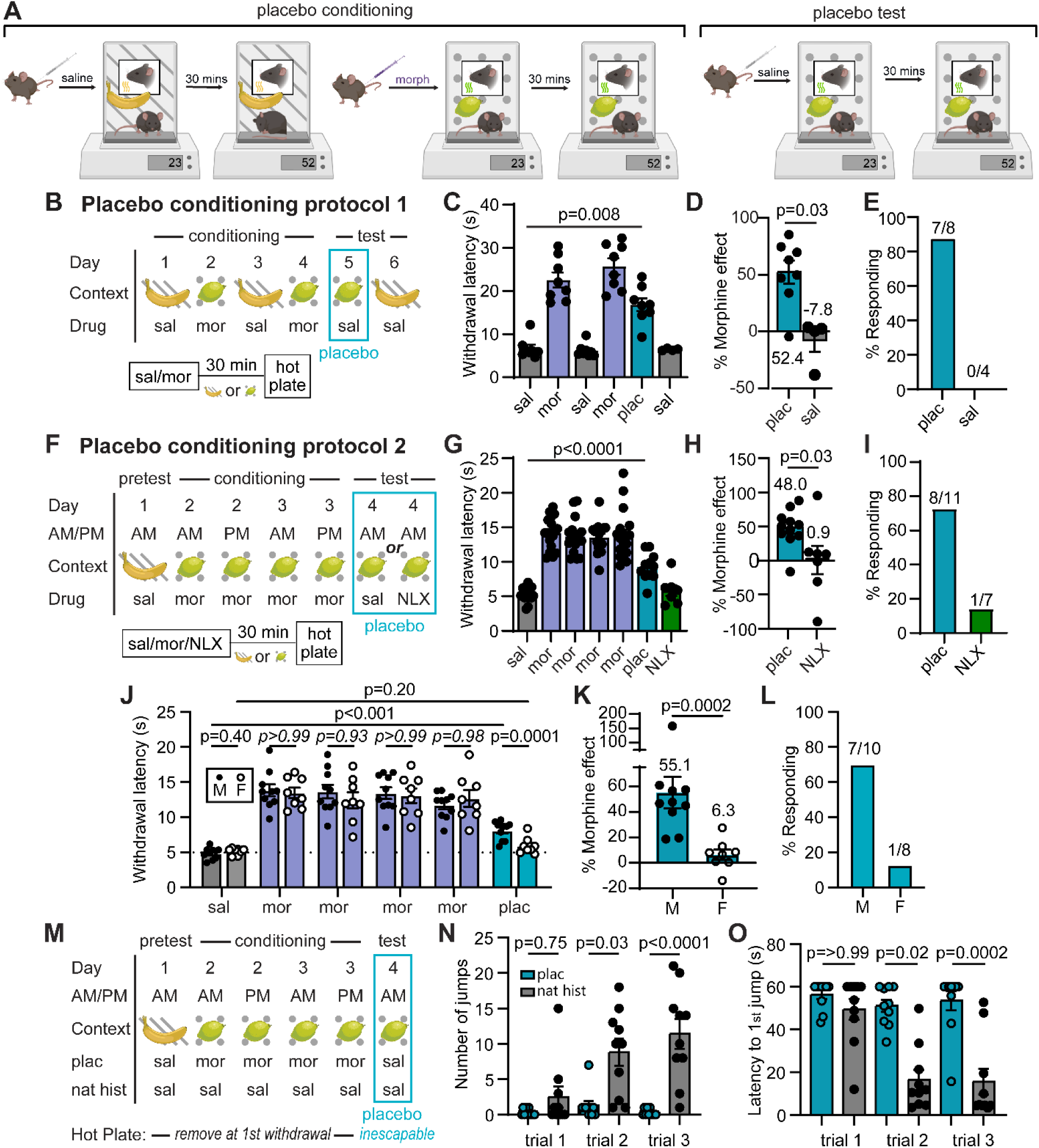
Morphine conditioning produces endogenous opioid-dependent placebo analgesia in male mice. (**A**) Illustration of placebo protocols based on contextual conditioning in chambers distinguished by both visual and olfactory cues. (**B**) Schematic of placebo protocol 1. (**C**) Hot plate paw withdrawal latencies before, during, and after placebo conditioning with morphine (n=8 mice, only 4 of which were tested on day 6, RM mixed effects model with Dunnett’s post-hoc, F(2.39,14.82)=40.58, p=0.008). (**D**) Placebo effect quantified as % of the response to the previous dose of morphine (Mann-Whitney test). (**E**) Placebo response quantified as the % of subjects exhibiting antinociception. (**F**) Schematic of placebo protocol 2, including naloxone administration on test day. (**G**) Same as (**C**), but in response to placebo protocol 2 (n=18 mice total, placebo: n=11, naloxone: n=7, RM mixed-effects model with Dunnett’s post-hoc, F(6,84)=50.85, p<0.0001). (**H**) Same as (**D**) (unpaired t-test). (**I**) Same as (**E**). (**J**) Same as (**G**) but comparing male and female mice (male: n=10, female: n=8, saline and placebo comparisons were made with a two-way ANOVA, saline/placebo vs. sex interaction: F(1,16)=17.04, p=0.008, uncorrected Fisher’s LSD post-hoc; morphine M/F comparisons were made using a one-way ANOVA with Sidak’s post-hoc (p-values italicized), F(11,96)=21.08, p<0.0001). (**K**) Same as (**D**). (**L**) Same as (**E**). (**M**) Schematic of placebo protocol 2 adapted to measure escape behaviors by transitioning to the inescapable hot plate on placebo test day. (**N**) The number of jumps across trials (placebo: n=10, natural history: n=10, Kruskal-Wallis test, H=35.05, p<0.0001 with Dunn’s post-hoc). (**O**) Latency to first jump across trials (Kruskal-Wallis test, H=34.71, p<0.0001 with Dunn’s post-hoc).

Protocol 2, which includes two morphine injections per day for two consecutive days and omits the saline context (**Figure 1F**), produced a placebo response similar to protocol 1. Consistent with a critical role for endogenous opioids^4,13,14^, treatment with naloxone (NLX, 3 mg/kg *i.p.*) on test day completely abolished the placebo effect (**Figure 1G-I**). Notably, the placebo response was not as robust in female mice, despite morphine producing similar analgesia during conditioning (**Figure 1J-L**), which is consistent with findings in healthy human subjects^28^. As a result, subsequent mechanistic studies were performed exclusively in male mice.

To determine if our placebo protocol reduces affective and/or emotional-motivational components of pain, we measured escape behaviors using the inescapable hot plate assay. After conditioning for the suppression of paw withdrawals, which reflect the sensory-discriminative component of pain (nociception), the inescapable hot plate assay was evaluated on placebo test day only. A natural history group administered saline instead of morphine was used as a control (**Figure 1M**). Strikingly, in addition to the increased paw withdrawal latency (**Extended Figure 1A-B**), mice in the placebo group exhibited reduced jumping behavior (**Figure 1N**) and an increased latency to the first jump (**Figure 1O**) in comparison to the natural history control mice. This was particularly apparent on trials 2 and 3, wherein control mice sensitized to the assay by increase their jumping behavior over subsequent trials, whereas the morphine-conditioned placebo group did not. Notably, the initial paw withdrawal latency was stable across trials in both groups (**Extended Figure 1C**). These results suggest that conditioning for the suppression of sensory-driven paw withdrawals also suppresses affective and/or emotional-motivational components of pain, and therefore produces analgesia, in addition to antinociception. Furthermore, because the suppression of escape-like jumping behavior was not included in the conditioning process, this form of morphine-conditioned placebo analgesia does not simply reflect a conditioned motor response.

To ask if the observed placebo effect is simply a consequence of non-specific, context-evoked endogenous opioid release^25,29,30^, we employed an unpaired variation of protocol 1 that decoupled the association between morphine context and pain suppression (**Extended Figure 1D-G**). Mice were tested on the hot plate prior to morphine administration and subsequent contextual conditioning so that they learned to associate the context with morphine, but never experienced analgesia on the hot plate. On test day, mice were administered saline in the morphine context prior to the hot plate test.

Consistent with the placebo effect being driven by a pain-predictive process, the unpaired conditioning protocol did not produce antinociception. Furthermore, using protocol 2, saline injection on placebo test day did not trigger morphine-associated behaviors such as locomotor activation or Straub tail (**Extended Figure 1H**), suggesting that saline injection during the placebo test does not evoke a general opioid-like behavioral response. We also found that decreasing the wait time on placebo test day between the saline injection and the hot plate test from 30 min to 10 min abolished the placebo effect (**Extended Fig. 1I-L**). Repeated testing caused the placebo response to extinguish with a time constant of 0.8 sessions (3 trials per session), and reconditioning with two morphine exposures reinstated the placebo effect to its initial magnitude (**Extended Figure 1M-P**). Together, these findings indicate that morphine-conditioned placebo analgesia relies on contextual and temporal discrimination to generate expectations that shape both sensory and affective behavioral responses to a noxious sensory stimulus.

### Placebo analgesia relies on the descending pain modulatory system

We set out to test the long-standing yet controversial hypothesis that opioid drugs and placebo analgesia both rely on activation of the DPMS. A central node in the DPMS is the ventrolateral PAG (vlPAG), which attenuates the processing of nociceptive sensory information in the spinal cord through its outputs to the rostroventromedial medulla (RVM) and the locus coeruleus^31–35^. Although it has not been directly demonstrated, vlPAG→RVM neurons are thought to be activated by opioids via a disinhibitory mechanism^36,37^. Prior studies have established that activation of glutamatergic vlPAG neurons produces analgesia^38,39^. We recently reported that ∼85% of vlPAG→RVM neurons are glutamatergic and that activity in vlPAG*^vGlut2-Cre^* neurons is required for systemic morphine antinociception on the hot plate via spinal opioidergic and noradrenergic signaling^35^.

To determine if the DPMS is engaged during morphine-conditioned placebo antinociception, we recorded hot plate-evoked Ca^2+^ activity from vlPAG→RVM projection neurons using fiber photometry before, during, and after conditioning with morphine using placebo protocol 2. Bilateral injection of AAVretro-Cre^40^ in the RVM and unilateral Cre-dependent jGCaMP8s^41^ in vlPAG provided access to vlPAG→RVM projection neurons (**Figure 2A**). Intriguingly, we observed the initiation of Ca^2+^ activity in vlPAG→RVM neurons during the process of transferring the mouse to the hot plate (**Figure 2B**), which may reflect the vlPAG’s role in threat processing^42^. It subsequently remained elevated during exposure to the noxious stimulus. Consistent with opioid-driven disinhibition of vlPAG→RVM neurons, this hot plate-evoked Ca^2+^ activity was enhanced by morphine (10 mg/kg, *i.p.*) and during the placebo test (**Figure 2C**).

**Figure 2:**
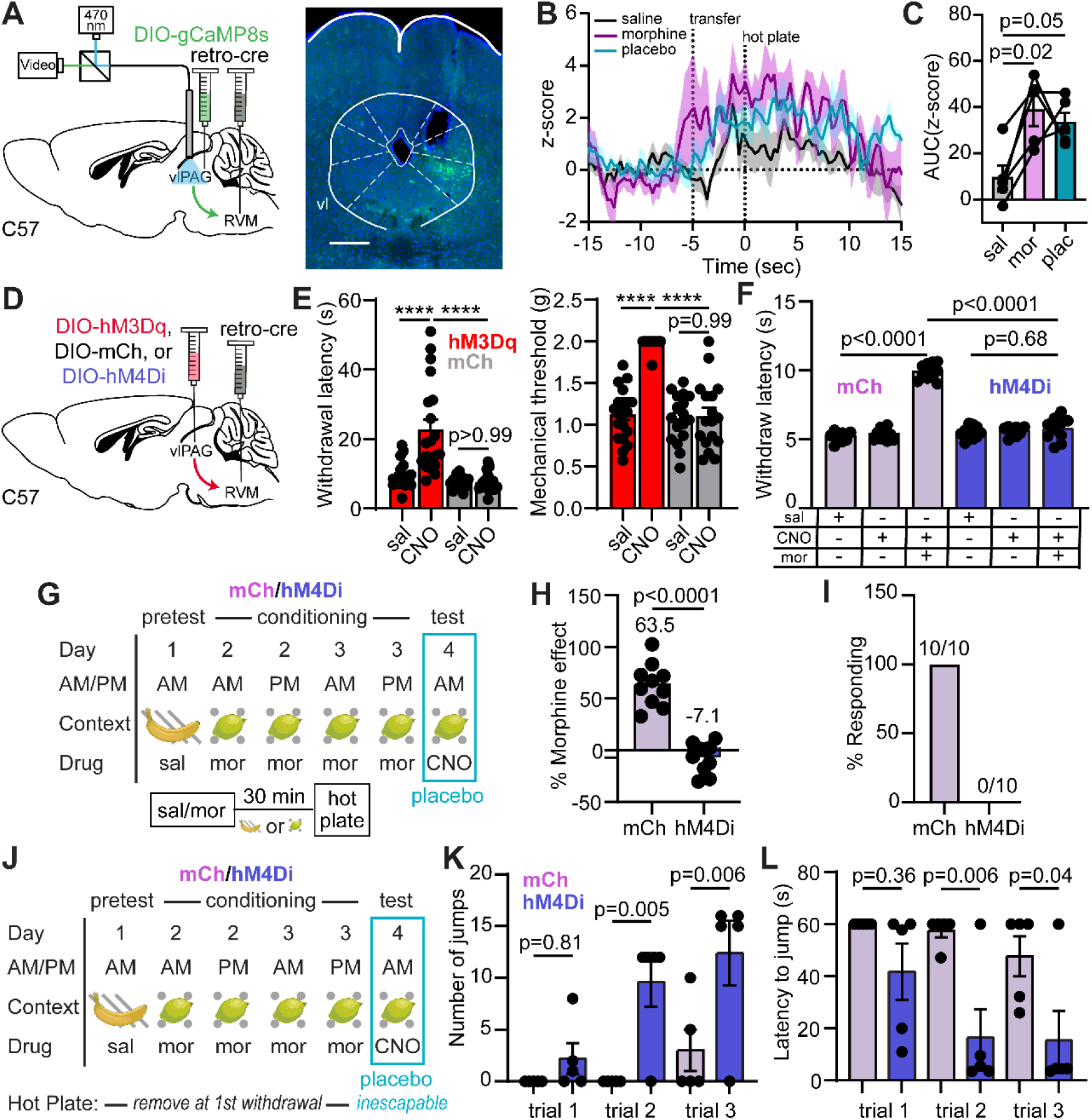
Pain modulatory vlPAG→RVM neurons support morphine analgesia and morphine-conditioned placebo analgesia. (**A**) Schematic of the retrograde viral injection approach for fiber photometry recordings from vlPAG→RVM neurons (left) and example of jGCaMP8s expression under the implanted optical fiber (right, scale bar = 0.5 mm). (**B**) Average z-score of Ca^2+^ activity in vlPAG→RVM neurons upon exposure to the hot plate over the course of placebo conditioning (protocol 2, n=5). (**C**) Quantification of the Ca^2+^ activity shown in (**B**) (integration window: -5 to 15 sec, RM one-way ANOVA with Holm-Sidak’s post-hoc, F(1.69,6.75)=7.37, p=0.022). (**D**) Schematic of the retrograde viral approach for DREADD expression in vlPAG→RVM neurons (examples of expression shown in **Extended Figures 2A-B**). (**E**) Paw withdrawal latencies on the hot plate (left) and mechanical thresholds to stimulation with von Frey fibers (right) upon chemogenetic activation of vlPAG→RVM neurons with hM3Dq vs. mCherry control (male and female mice combined, hM3Dq: n=20, mCh: n=17, HP: One-way ANOVA with Sidak’s post-hoc, F(3,70)=19.64, p<0.0001; VF: One-way ANOVA with Sidak’s post-hoc, F(3,70)=43.36, p<0.0001), ****=p<0.0001. (**F**) Hot plate paw withdrawal latencies in response to saline or morphine (5 mg/kg) administration with chemogenetic inactivation of vlPAG→RVM neurons with hM4Di vs. mCherry control (mCh: n=10, hM4Di: n=10, One-way ANOVA with Sidak’s post-hoc, F(5,54)=115.6, p<0.0001). (**G**) Schematic of placebo protocol 2 indicating chemogenetic silencing with CNO administration on placebo test day. (**H**) Placebo effect quantified as % of the response to the previous dose of morphine upon chemogenetic silencing of vlPAG→RVM neurons (mCh: n=10, hM4Di: n=10, unpaired t-test). (**I**) Placebo response in (**H**) quantified as the % of subjects exhibiting antinociception. (**J**) As in (**G**), but to measure escape behavior on placebo test day. (**K**) The number of jumps across trials upon chemogenetic silencing of vlPAG→RVM neurons (mCh: n=5, hM4Di: n=5, One-way ANOVA with Sidak’s post-hoc, F(5,24)=7.56, p=0.0002). (**L**) Latency to first jump across trials (One-way ANOVA with Sidak’s post-hoc, F(5,24)=5.42, p=0.002).

Consistent with vlPAG→RVM neurons comprising a key node in the DPMS, their bilateral chemogenetic activation with the excitatory DREADD hM3Dq^43^ (3 mg/kg CNO, *i.p.*) produced strong antinociception on the hot plate (**Figure 2D-E**, **Extended Figure 2A**). Furthermore, chemogenetic silencing of vlPAG→RVM neurons with hM4Di prevented morphine (5 mg/kg, *i.p.*) antinociception (**Figure 2F**, **Extended Figure 2B-C**), similar to vlPAG*^vGlut2-Cre^* neurons^35^. Strikingly, chemogenetic silencing of either vlPAG→RVM neurons (**Figure 2G-I**, **Extended Figure 2D**) or vlPAG*^vGlut2-Cre^* neurons (**Extended Figure 2E-J**) on placebo test day completely blocked placebo antinociception. It also reversed the placebo-driven suppression of unconditioned escape-like jumping behavior (**Figure 2J-L**). These results establish that morphine-conditioned placebo analgesia, similar to morphine analgesia itself, requires engagement of the DPMS through the vlPAG.

### Neural pathways for top-down pain modulation

Although multiple cortical structures are implicated in placebo pain relief, it is not clear how the PAG receives top-down information that drives the DPMS. We established a viral approach to access PAG-projecting cortical neurons using retrograde AAV (AAVretro). Bilateral injection of AAVretro encoding tdTomato (AAVretro-tdTom) in the vlPAG led to dense labeling of cortical neurons located throughout the rostro-caudal axis of anterior cingulate cortex (ACC), the infralimbic (IL) and prelimbic (PrL) cortices (collectively referred to here as the medial PFC (mPFC)), secondary motor cortex (M2), and the anterior insula (AI) (**Figure 3A-B**).

**Figure 3:**
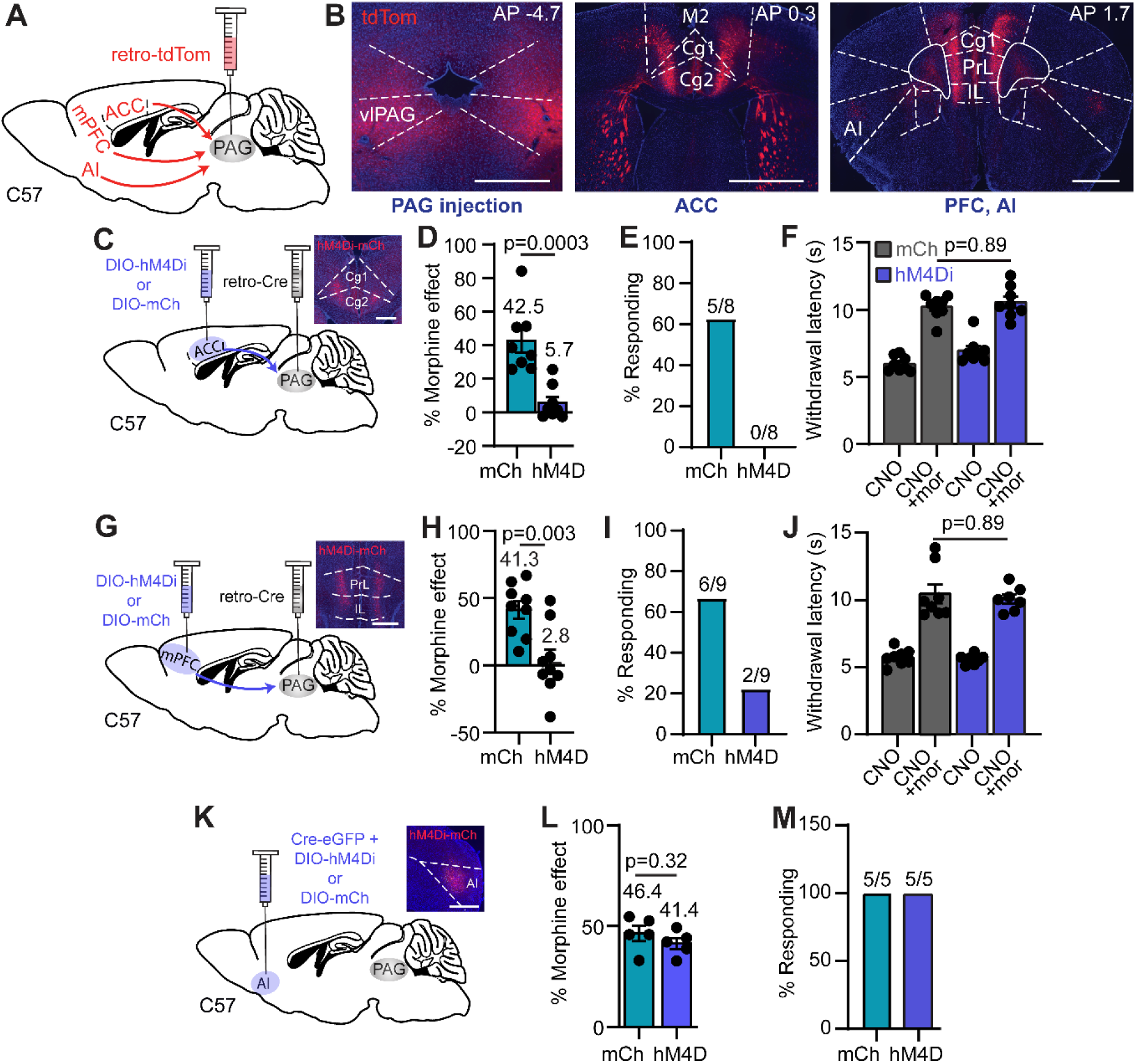
Cortical input to the PAG is critical for placebo analgesia. (**A**) Schematic of AAVretro tdTom injection into vlPAG for retrograde labeling of cortical inputs. (**B**) Example images of the bilateral vlPAG injection sites (left, scale bar 0.5 mm) and labeled input neurons in the ACC (middle, scale bar 1 mm) and mPFC (right, scale bar 1 mm). (**C**) Schematic of the retrograde viral injection approach for chemogenetic silencing of ACC→PAG neurons. Inset: representative image of hM4Di-mCh expression in ACC (scale bar 0.5 mm). (**D**) Placebo effect quantified as % of the response to the previous dose of morphine upon chemogenetic silencing of ACC→PAG neurons (mCh: n=8, hM4Di: n=8, unpaired t-test). (**E**) Placebo response in (**D**) quantified as the % of subjects exhibiting antinociception. (**F**) Hot plate paw withdrawal latencies in response to CNO (3 mg/kg, *i.p.*) or CNO + morphine (5 mg/kg) administration with chemogenetic inactivation of ACC→PAG neurons with hM4Di vs. mCherry control (mCh: n=8, hM4Di: n=8, RM one-way ANOVA with Sidak’s post-hoc, F(3,28)=47.74, p<0.0001). (**G**) Same as **C** but for mPFC→PAG neurons. (**H**) Same as **D** but for mPFC→PAG neurons (mCh: n=9, hM4Di: n=9, unpaired t-test). (**I**) Same as **E** but for mPFC→PAG neurons. (**J**) Same as **F** but for mPFC→PAG neurons (mCh: n=8, hM4Di: n=7, RM one-way ANOVA with Sidak’s post-hoc, F(3,26)=39.90, p<0.0001). (**K**) Same as **C** for the PAG-projecting region of AI (scale bar 1 mm). (**L**) Same as **D** but for the PAG-projecting region of AI (mCh: n=5, hM4Di: n=5, unpaired t-test). (**M**) Same as **E** but for the PAG-projecting region of AI.

To assess the possibility that cortical input to the PAG contributes to the placebo effect, we chemogenetically silenced PAG-projecting neurons located in the ACC, mPFC, and AI during the placebo test. PAG-projecting neurons in either the ACC or mPFC were labeled by bilaterally injecting AAVretro-Cre in the vlPAG, along with either Cre-dependent hM4Di or mCherry in cortex, whereas the PAG-projecting region of AI was directly transduced with a mixture of Cre and hM4Di. Consistent with causal roles for the ACC and mPFC, placebo antinociception was blocked by inactivation of PAG-projecting neurons in the ACC and mPFC (**Figure 3C-E**, **Extended Figure 3A** and **Figure 3G-I**, **Extended Figure 3B**, respectively). Notably, bilateral chemogenetic silencing of ACC-to-PAG or mPFC-to-PAG neurons did not alter morphine antinociception (5 mg/kg, *i.p.*, **Figure 3F and 3J**). In contrast to the ACC and mPFC, inactivation of the PAG-projecting region of the AI had no effect on placebo antinociception (**Figure 3K-M**, **Extended Figure 3C**), consistent with an evaluative role for the AI in placebo analgesia^9,15,44^. To determine if these cortical inputs to the PAG are sufficient to drive antinociception, which has been observed for the mPFC→PAG pathway after injury^45^, we simultaneously activated both ACC→PAG and mPFC→PAG neurons bilaterally with hM3Dq. However, this failed to alter nociception using either the hot plate or von Frey assays (**Extended Figure 3D-E**).

Together, these results indicate that cortical inputs to the PAG from the ACC and mPFC contribute causally to placebo pain relief. In contrast, cortical input to the PAG is not a key factor in the analgesia produced by opioid drugs. Yet, cortical input from the ACC and mPFC is unable to drive descending pain modulation through the PAG on its own, suggesting that an additional process is critical for triggering the DPMS to produce placebo analgesia.

### Endogenous opioid signaling in placebo analgesia

Like most forms of placebo pain relief, morphine conditioned placebo relies on endogenous opioid signaling. We hypothesized that this may occur in the vlPAG, as local administration of opioid drugs in the vlPAG activates the DPMS to produces strong antinociception^46^. Although PET imaging studies in humans suggest that endogenous opioid peptides are released in the PAG during placebo, a causal role for local opioid signaling has not been established. Furthermore, the temporal dynamics of opioid release during placebo remain undefined.

To determine if morphine-conditioned placebo analgesia involves endogenous opioid peptide release in the vlPAG, we used fiber photometry to measure fluorescence from the genetically encoded opioid sensor δLight^47^. By transducing *Oprm1-Cre* mice^48^, sensor expression was restricted to vlPAG*^Oprm1-Cre^* neurons, which are the most likely targets of endogenous opioid signaling (**Figure 4A**). Prior to conditioning, placement on the hot plate caused a sustained reduction in sensor fluorescence, which could reflect either an acute loss of opioid tone or, more likely, sensor quenching due to activity-dependent acidification near the plasma membrane inner leaflet in sensor-expressing neurons^49^ (**Figure 4B-C**). In contrast, on placebo trials, sensor fluorescence rapidly increased over the first 5 seconds of noxious stimulus exposure and remained elevated for at least 10 seconds thereafter. This finding suggests that placebo conditioning increases noxious stimulus-driven endogenous opioid peptide signaling in the vlPAG.

**Figure 4.**
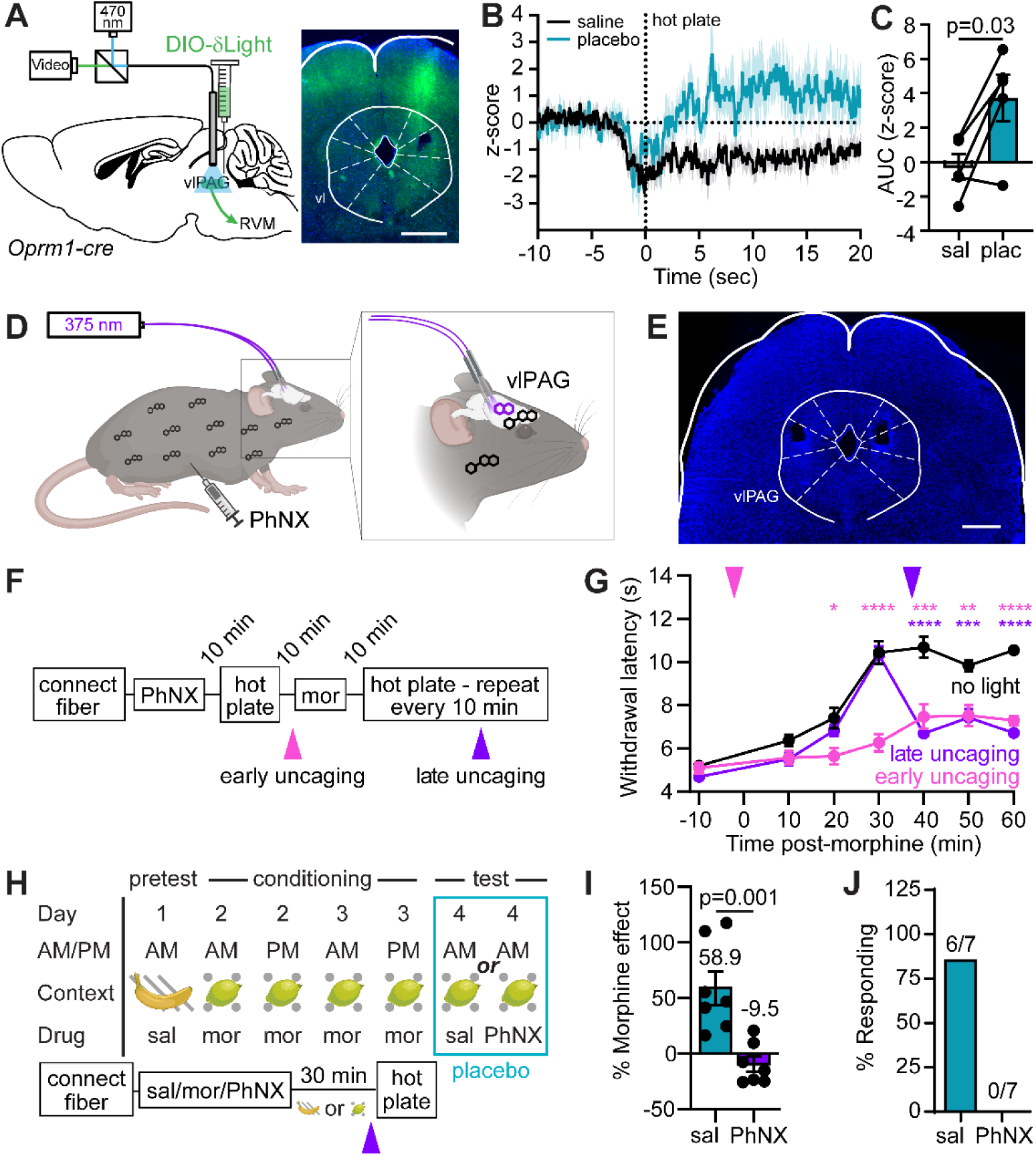
vlPAG opioid signaling underlies morphine antinociception and placebo analgesia. (**A**) Schematic of the viral injection approach for fiber photometry recordings of endogenous opioid peptide signaling from vlPAG*^Oprm1-Cre^* neurons (left) and example of δLight expression under the implanted optical fiber (right, scale bar 0.5 mm). (**B**) Average z-score of opioid peptide activity in vlPAG*^Oprm1-Cre^* neurons upon exposure to the hot plate before and after placebo conditioning (protocol 2, n=5). (**C**) Quantification of the opioid activity shown in (**B**) (integration window: 3 to 8 sec, paired t-test). (**D**) Schematic the configuration used for bilateral *in vivo* photorelease of naloxone from the caged opioid antagonist PhNX in the vlPAG. (**E**) Example of bilateral optical fiber implantation over the vlPAG (scale bar 1 mm). (**F**) Timeline for temporally-defined *in vivo* naloxone photorelease during morphine antinociception. (**G**) Time-course of hot plate withdrawal latencies in response to morphine (5 mg/kg) and PhNX (30 mg/kg) administration with light applied in the vlPAG as indicated (no light: n=11, early: n=10, late: n=10, Two-way repeated measures ANOVA with Dunnett’s post-hoc, Time x Condition F(12,168)=13.54, p<0.001, asterisks are color coded for early and late uncaging compared to the no light condition, *=p<0.05, **=p<0.01, ***=p<0.001, ****=p<0.0001). (**H**) Schematic of placebo protocol 2 modified for *in vivo* PhNX photoactivation 3 minutes prior to hot plate presentation on placebo test day. (**I**) Placebo effect quantified as % of the response to the previous dose of morphine upon naloxone photorelease in the vlPAG vs. saline control (saline: n=7, PhNX: n=7, unpaired t-test). (**J**) Placebo response in (**I**) quantified as the % of subjects exhibiting antinociception.

To determine if vlPAG opioid signaling is necessary for placebo antinociception, we implemented a novel photopharmacological approach involving bilateral photoactivation of PhNX, a photo-caged, blood brain barrier-permeable derivative of the opioid antagonist naloxone^50^ (**Figure 4D-E**). This approach was imperative, as bilateral cannulation lesions much of the PAG, and the excessive animal handling required for drug infusion interferes with placebo conditioning. Further confirming PhNX’s inactivity at opioid receptors *in vivo*, systemic PhNX administration (30 mg/kg, *i.p.*) did not attenuate morphine (5 mg/kg, *i.p.*) antinociception in the hot plate assay in the absence of illumination (**Extended Figure 4A-B**). We first determined that bilateral vlPAG illumination with brief 375 nm light flashes (10 x 200 ms, 1 Hz, 22.5 mW) either 2 minutes before (early) or 37 minutes after (late) administration of morphine (5 mg/kg *i.p.*) (**Figure 4F**) strongly attenuated the resulting antinociception (**Figure 4G**). Notably, early and late naloxone photorelease produced similar effects, indicating that opioid signaling in vlPAG is necessary for maximal morphine antinociception.

Next, we bilaterally photoreleased naloxone on placebo test day (**Figure 4H**). Importantly, PhNX had no effect on placebo in the absence of illumination (**Extended Figure 4C-E**). To limit our manipulation to a time window involving the noxious stimulus, we photoactivated PhNX (30 mg/kg *i.p.*) 27 minutes after placing mice in the morphine-paired chamber, just 3 minutes prior to hot plate exposure. In comparison to treatment with light and saline, PhNX photoactivation in vlPAG completely blocked the placebo effect (**Figure 4I-J**). These results establish the vlPAG as a critical site of endogenous opioid signaling during the pain-processing phase of placebo pain relief.

## Discussion

The lack of validated preclinical models of placebo analgesia has hindered studies into the neural mechanisms of placebo pain relief. Our study establishes the relevance of morphine-conditioning in mice as a means of generating pain-related expectations to produce a placebo effect that shares multiple features with placebo analgesia in humans. Strikingly, we found that conditioning for antinociception suppresses emotional/motivational components of pain-related behavior. To the best of our knowledge, this is the first time that conditioning in rodents has been demonstrated to produce a placebo effect that extends beyond the conditioning stimuli. This is significant because humans participating in clinical trials exhibit strong placebo effects that result, at least in part, from the generalization of prior experiences with other therapeutics^5^.

The relevance of the DPMS in placebo analgesia has been a subject of continuous debate^20–23^. Our study establishes a central role for the DPMS in placebo analgesia and identifies that ACC and mPFC projections to the PAG are critical neural pathways through which prefrontal cortical circuits implement top-down control over pain processing. Whereas morphine antinociception and morphine-conditioned placebo antinociception both rely on neural activity and opioid signaling in the vlPAG, cortical input to the PAG is uniquely involved in placebo pain relief. By revealing a critical role for cortical neurons that project directly to the PAG, our work provides a neural circuit basis for prior correlational observations implicating the PFC^51,52^, along with metrics of PFC-PAG^12^ and ACC-PAG connectivity^11,12,17^ as being important for placebo analgesia.

A recent study in mice produced a placebo effect by optogenetically stimulating anesthesia-activated neurons in the central amygdala during conditioning^53^. That placebo effect did not rely on re-activation of the neurons used for conditioning, which stands in contrast to the necessity of vlPAG→RVM neurons in both morphine conditioning and placebo analgesia in our study. Another study reported that ACC projections to the pontine nucleus were critical for a form of conditioned placebo that does not involve opioid drugs, but that work did not investigate the PAG^54^. Whether or not these related, yet distinct forms of placebo pain relief rely on shared neural mechanisms remains to be determined.

It has been suggested that endogenous opioid release in the PAG is not a primary factor in placebo analgesia^55,56^. Instead, our results using a novel opioid sensor^47^ and a novel photopharmacological method^50^ establish causal roles for the DPMS and endogenous opioid signaling in the vlPAG. Our results point to a mechanism of pain relief in which noxious stimuli acutely drive endogenous opioid peptide release in the vlPAG to activate the DPMS, which, in turn, suppresses the incoming noxious sensory information in the spinal cord on the timescale of seconds. This negative feedback loop is enhanced by placebo conditioning through a currently unknown mechanism. Altogether, our findings support a model in which the PAG integrates top-down information pertaining to expectations about noxious stimuli with local endogenous opioid signaling to activate the DPMS, thus producing placebo pain relief.

Our placebo model relies on contextual conditioning with morphine, raising the question of whether the resulting analgesia can be attributed to expectations about pain, as opposed to pre-cognitive learned associations. Careful psychopharmacological studies in humans have shown that expectation-dependent placebo analgesia is fully opioid dependent, whereas extensive conditioning produces a firm, expectation-independent placebo component that depends on endocannabinoids^4,57^. The opioid dependence of our placebo effect is consistent with an expectation-dependent process.

Because the unpaired conditioning protocol failed to produce a placebo response, we conclude that our placebo protocol likely involves acute expectations about the aversive qualities of the noxious stimulus. Thus, our findings are congruent with a predictive coding model of placebo analgesia in which the vlPAG tunes ascending sensory signals to more closely match predictions about noxious sensory stimuli via activation of the DPMS^3,58^.

Additional work is required to unravel the underlying PAG microcircuit, including the mechanisms governing opioid peptide release, both of which will be important for determining if the circuit can indeed support predictive coding^3,59^. The lack of placebo observed in female mice may relate to well documented sex-differences in the anatomy and neurochemistry of the vlPAG→RVM circuit^60^. Although a similar trend has been documented in uninjured humans^28^, such a sex difference has not been observed in patients suffering from chronic pain. Further evaluation of these circuits in preclinical chronic models will be required to understand how the circuit might be transformed by injury, and how the mechanisms uncovered here might be harnessed to optimize placebo effects in the clinic, which could involve the use of cognitive behavioral therapy to engage or reshape the underlying neural pathways^61^.

## Materials and Methods

### Animals

All procedures were approved by the UCSD Institutional Animal Care and Use Committee and guidelines of the National Institute of Health. Mice were housed 1–5 per cage and maintained on a 12-hour reversed light/dark cycle in a temperature-controlled environment with *ad libitum* access to food and water. Experiments were performed under red lighting during the dark period using either C57Bl/6J mice (Jackson Laboratory, stock #664, males, aged 8–60 weeks), *Oprm1-Cre* knock-in mice (Jackson Laboratory, stock # 038053), or *vGlut2-*IRES-Cre knock-in mice (Jackson Laboratory, stock # 028863, aged 8–60 weeks).

### Drugs

Clozapine-N-oxide (CNO) was purchased from Hello Bio, (HB6149). Morphine (mor) was purchased from Spectrum Chemicals (M1167). Saline solution was purchased from Teknova (S5819). Naloxone hydrochloride (NLX) was purchased from Tocris (0599). Photoactivatable Naloxone (PhNX) was synthesized according to the published procedure^50^ and purified to greater than 99.9% purity. Morphine was obtained from Spectrum (Cat:M1167).

### Viral constructs

The following viruses were obtained from Addgene: AAVretro-pgk-Cre (Addgene 24593, titer 1×10^13^ GC/ml), AAVretro-hSyn-Cre (Addgene 105553-AAVrg, titer 2.1×10^13^ GC/ml), AAVretro-CAG-tdTom (Addgene 59462, titer 1.2×10^12^ GC/ml), AAV8-hsyn-DIO-hM4Di-mCherry (Addgene 44362, titer 2.3×10^12^ GC/ml), AAV8-hsyn-DIO-hM3Dq-mCherry (Addgene 44361, titer 2.1×10^12^ GC/ml), AAV8-hsyn-DIO-mCherry (Addgene 50459, titer 3.5×10^12^ GC/ml), AAV1-AAV-syn-FLEX-jGCaMP8s-WPRE (162377, titer 2.3×10^12^ GC/ml). AAV-DJ-CBA-Cre-eGFP (Addgene 49056, titer 4.3×10^12^ GC/ml) was produced in house using plasmid obtained from Addgene and purified by ultracentrifugation with an iodixanol gradient. Titer was determined by qPCR. AAV9-Syn-tTA-TRE-DIO-δLight (titer 4.93×10^12^ GC/ml) was obtained from UNC Neurotools.

### Surgery

Before surgery, mice were deeply anesthetized by induction at 5% isoflurane, after which anesthesia was maintained by 2% isoflurane (SomnoSuite, Kent Scientific). After mice were placed in a stereotaxic frame (David Kopf Instruments), a midline incision was made through the scalp following fur removal and site preparation by alternating povidone-iodine and 70% isopropyl alcohol. 150-250 nl of virus was injected at a rate of 100 nl/min. The following coordinates were used for viral injection and optical fiber implantation, with DV indicating the distance ventral to skull. vlPAG: (angle ±10°) AP−4.60 mm, ML ±0.32 mm, DV 2.72 mm; RVM: (bilateral, angle -10°) AP -7.00 mm, ML +0.26 and -0.67 mm, DV +6.30 and +6.26 mm; ACC: AP 1.20 mm, ML ±0.30 mm, DV 2.52 mm; mPFC: AP 1.98 mm, ML ±0.30 mm, DV 2.20 mm; AI: AP 2.0, ML -2.50 mm and +2.20 mm, DV 3.20 mm. For all surgeries, mice were administered 5 mg/kg ketoprofen (MWI Veterinary Supply) before the end of surgery and 24 hours later and monitored for recovery for 5 days. Mice were given at least one week to recover from cannula or optical implant surgeries before behavior or fiber photometry recordings.

### Chemogenetic manipulation

In all experiments, experimenters were blind to the AAV with which experimental animals were transduced (DREADD or mCherry control). Mice were underwent behavioral testing 6 weeks after viral injection. The dose of CNO (3 mg/kg) was chosen based on published studies and was confirmed to induce robust cFOS expression in hM3Dq-labeled neurons in our hands (data not shown). In morphine analgesia experiments, CNO was injected *i.p.* 60 min before beginning behavioral test followed by *i.p.* injection of morphine (5 mg/kg, 10 mg/kg and 20 mg/kg) 30 min later.

### Pain behavior assessment

Thermal nociception was assayed using the hot plate test. The test was performed to evaluate heat sensitivity thresholds, measuring latency of hind paw withdrawal, hind paw licking, or jumping, after being manually placed on the heat source, which was calibrated weekly (ThermoFisher Scientific Hot Plate #SP88857100). Mice were tested on hot plate 3 times, every 5 minutes, and measurements were averaged to obtain a threshold value for each animal. The cutoff time of 60 sec to avoid tissue damage was never reached. In experiments involving vlPAG-mediated analgesia (**Figure 2E**), both male and female mice were used in equal numbers and the data were combined, as no sex difference was observed. The experimenter was blind to either drug identity or virus identity.

### Placebo analgesia

Other than in **Figure 1J-K**, placebo experiments were conducted in male mice, as female mice did not exhibit robust a placebo effect. Most experiments were conducted with two separate cohorts of 8-10 mice each. Each cohort contained control and experimental subjects in roughly equal numbers, and data from both cohorts were pooled.

Two placebo protocols were employed. Placebo protocol 1 is similar to the protocol of Guo et al.^16^. Mice were conditioned for 4 consecutive days, via injection with either saline or morphine (10 mg/kg, *i.p.*) followed immediately by placement in a cylindrical conditioning chamber (4.5” diameter, 8” height) with distinct contexts composed of a combination of visual cues (diagonal stripes or dots) and olfactory cues (limonene (citrus) or isoamyl acetate (banana)). Of note, although *n*-pentyl acetate, another banana-like odorant, has been found to produce stress-induced analgesia in male mice^62^, isoamyl acetate did not alter hot plate paw withdrawals. The odorants were counterbalanced with the visual cues and presented by taping to the inner wall of the chamber a q-tip loaded with 1% of the odorant in mineral oil, such that a standing mouse could not reach the tip. Context pairings were randomized and balanced across tested cohorts. The floor of the waiting chamber was a room temperature hot plate.

After 30 minutes in the conditioning chamber, mice were transferred, along with their conditioning chamber, to an adjacent pre-heated hot plate, and the animal’s latency to respond to the noxious heat (52°C) was assessed. Each hot plate test was repeated 3 times with at 5 minute intervals and the latencies were averaged. On day 5, the mice were administered saline and then placed in the conditioning chamber associated with morphine.

Placebo protocol 2 omits the intermittent saline conditioning periods and uses a lower dose of morphine (10 mg/kg, *i.p.*) to minimize tolerance. On the pre-test day, mice were administered saline and withdrawal baseline threshold was measured for each animal.

Each hot plate test was repeated 3 times at 5 minute intervals and the latencies were averaged. Subsequently, for 2 consecutive days (conditioning days 1 & 2) mice were injected twice a day (am and pm) with morphine and then placed in the conditioning chamber. After 30 minutes, mice were transferred with their conditioning chamber to a pre-heated hot plate, and the animal’s latency to respond to the noxious heat (52°C) was assessed. On day 3 (test day), the mice were administered saline, CNO (mg/kg, *i.p.)*, naloxone (10 mg/kg, *i.p.)*, or PhNX (30 mg/kg, *i.p.)* and then placed in the conditioning chamber associated with morphine.

To calculate the percentage of morphine effect in the placebo experiments, we applied the following formulas:

Placebo protocol 1:

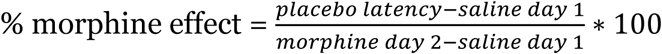

Placebo protocol 2:

9

On placebo test day, mice were considered responders if they exhibited withdrawal latencies >3 standard deviations greater than the mean on saline day 1 (pre-test), which includes the pooled data from up to 2 cohorts.

To measure affective and/or emotional-motivational components of pain during placebo analgesia, we measured the number of jumps and latency to first jump across three trials (5 minute intervals) on the hot plate during test day. A 60 second cut-off was used to prevent tissue damage to the paws.

### Fiber photometry

For fiber photometry recordings in vlPAG from vlPAG→RVM neurons, C57Bl/6J mice were injected bilaterally in the RVM with AAVretro-hSyn-Cre (150 nl per injection) and unilaterally in the vlPAG with a 1:1 mixture of AAV1-syn-FLEX-jGCaMP8s + AAVDJ-hsyn-DIO-mCherry (150 nl) and implanted unilaterally in the right vlPAG with a 200 µm diameter optical fiber. For recordings from ACC→PAG and mPFC→PAG neurons, C57Bl/6J mice were unilaterally injected in the right vlPAG with AAVretro-pgk-Cre (150 nl) and the right ACC or mPFC with a 1:1 mixture of AAV1-syn-FLEX-jGCaMP8s + AAVDJ-hsyn-DIO-mCherry (250 nl) and implanted in the ipsilateral right ACC or mPFC with a 200 µm diameter optical fiber. For recordings of δLight in vlPAG, *Oprm1-Cre* mice were unilaterally injected in the right vlPAG with AAV9-Syn-tTA-TRE-DIO-δLight (150 nl) and implanted in the vlPAG with a 200 µm diameter optical fiber. Recordings were made using a Neurophotometrics FP3001 fiber photometry system at a sampling rate of 30 Hz. A 470 nm LED was used to record GCaMP activity with a 560 nm LED for the mCherry control channel. For δLight recordings, a 470 nm LED was used to record δLight activity and 415 nm LED (isosbestic point) was used as control. Data analysis of fluorescence recordings was conducted using the pMAT open source photometry analysis package^63^ in MATLAB 2021a (Mathworks), which allows for the scaling of the red control channel to correct for movement and bleaching, and subsequently calculates peri-event time histograms of ΔF/F and z-score values. Other calculations, such as baseline correction, downsampling, and area under the curve analysis, were completed in Igor Pro (Wavemetrics). Fluorescence traces are presented in the figures as the mean ± SEM of single traces (hot plate trial 1) from all mice.

### *In vivo* drug uncaging

Ultraviolet light was delivered to the mouse brain through optical fibers (200 µm diameter, high-OH, 1.25 mm ceramic ferrule, Neurophotometrics, Ltd) implanted bilaterally over the vlPAG. Light from a 375 nm laser (Oxxius, LBX-375-400-HPE-PPA) was launched into two 200 µm diameter, high-OH fiber optic cables (Thorlabs) using a custom-built light path that included a 50/50 UV beamsplitter to direct the light into the two fiber optic cables. Light power was calibrated immediately prior to use to deliver 30 mW from the cable tip. Transmission through optical fibers prior to implantation was ∼75% such that estimated power delivery to the brain was ∼22.5 mW. Light pulses (10 x 200 ms, 1 Hz) were generated in response to experimenter-controlled switch by an Arduino UNO.

For PhNX uncaging during morphine analgesia, after being bilaterally connected to the fiber optic cables, mice were injected with PhNX (30 mg/kg, *i.p*) and a baseline thermal nociception was assayed using the hot plate test 10 minutes after acclimation. For early uncaging experiments, PhNX was uncaged 1 minute before injection of morphine (5 mg/kg, *i.p*). Subsequently, baseline thermal nociception was assayed using the hot plate every 10 minutes for a total of 60 minutes. For late uncaging experiments, PhNX was uncaged 37 minutes after morphine injection. For no-light control experiments, PhNX was never photoactivated. For this timecourse experiment, hot plate measurements were only taken once per time point.

For experiments involving uncaging during placebo analgesia, baseline thresholds were acquired, and mice were conditioned while bilaterally connected to the fiber optic cables but no light was applied. On placebo test day mice were injected with either saline or PhNX (30 mg/kg *i.p*) prior to placement in the morphine context. Light flashes were triggered 3 minutes before placing mice on the hot plate 3 times at a 5 minute interval.

### Histology

Brain tissue was fixed using 4% paraformaldehyde and sectioned on a freezing microtome (Leica). After mounting on glass coverslips in DAPI-containing medium (Vector Laboratories, #H1200), sections were imaged using a Keyence microscope (BZ-X710) and subsequently processed in Image J.

### Statistics

Throughout the manuscript, data are represented as the mean±SEM. Data were analyzed in GraphPad Prism. All datasets were evaluated for normality using the D’Agostino and Pearson test and the Shapiro-Wilk test. Normally-distributed datasets were analyzed using parametric statistics and non-normally-distributed data sets were analyzed using non-parametric statistics, taking into account repeated measures from individual subjects when appropriate. P values are presented in the figures. The number of experimental replicates and the statistical test used are presented in each figure legend. In some cases, when cohorts were split into multiple conditions (e.g. saline vs. naloxone), instead of a two-way ANOVA, the equivalent Mixed-effects model was used to allow for missing data. Not all statistically significant comparisons are indicated in the figures. Raw data, summary data, and statistical test details (including F and t values, and degrees of freedom) for each graph are available in the Source Data spreadsheet.

## Acknowledgements

The authors thank J. Isaacson, M. Smith, F. Zeidan, S. Han, E. Azim, T. Yaksh, B.K. Lim, S. Sternson, G. Pekkurnaz, and S.P. McClain for helpful discussions, B. Stepanyuk and E. Berg for AAV production, and E. Osgood for assistance with surgeries. Figures 1A and 4D were partially created with BioRender.com.

## Author contributions

G.L., S.T.L., and M.R.B. conceived the project. G.L., J.L., D.A.J., X.M., S.T.L., J.P., and J. C-W. designed the work and performed experiments. C.D. and L.T. generated, characterized, and provided δLight. G.L., J.L., and M.R.B. wrote the manuscript. All authors interpreted the data, contributed to data analysis, and discussed the manuscript.

## Materials & Correspondence

Correspondence and material requests should be addressed to Dr. Matthew Banghart. All data are available in the Source Data file.

---

**Extended Figure 1:**
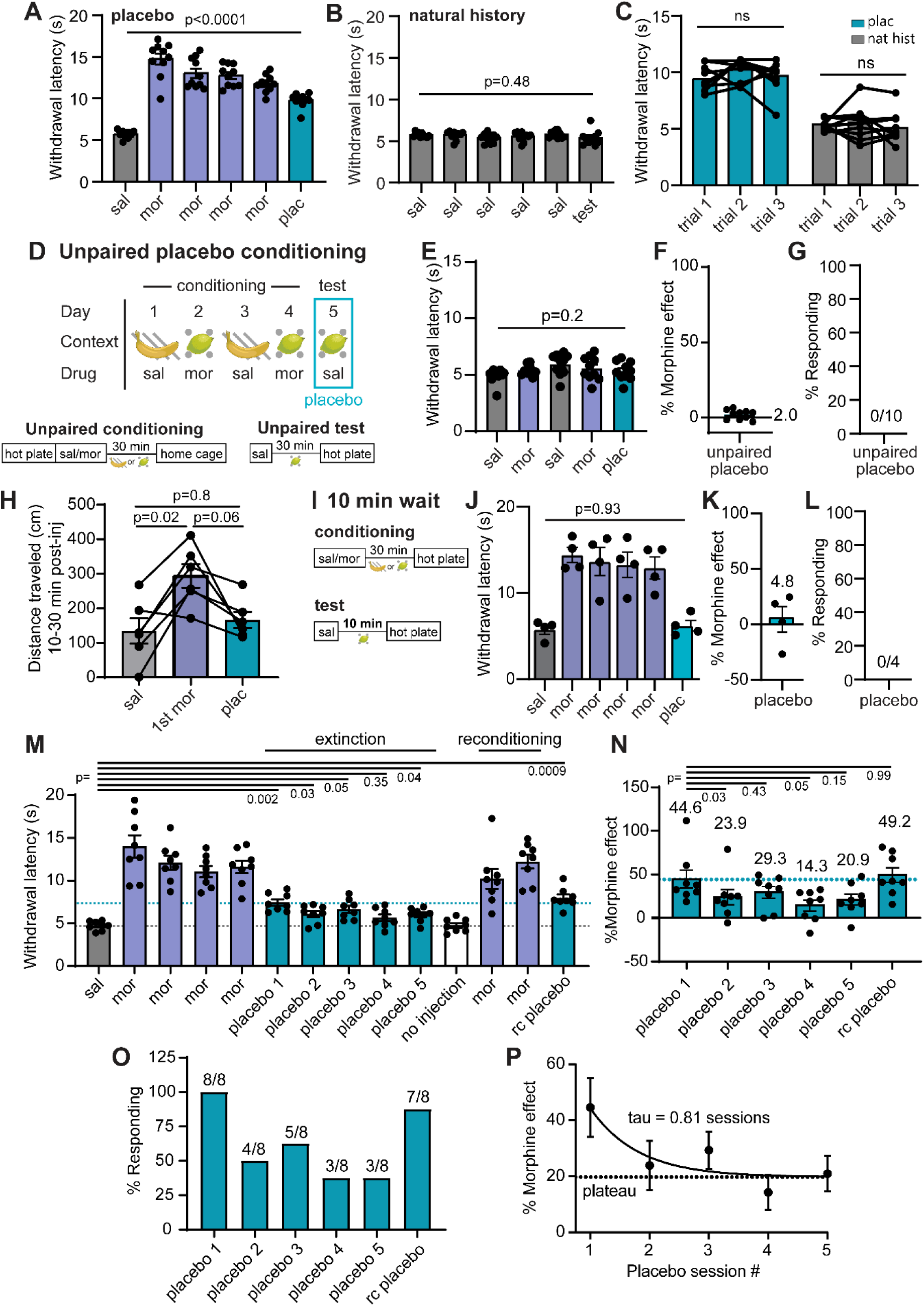
Further characterization of morphine-conditioned placebo analgesia. (**A**) Hot plate paw withdrawal latencies before, during, and after placebo conditioning with morphine for the placebo group shown in Figure 1M-O (n=10, RM one-way ANOVA with Dunnett’s post-hoc, F(2.41,21.64)=71.40, p<0.001). (**B**) Same as A, but for the natural history group (n=10, RM one-way ANOVA with Dunnett’s post-hoc, F(2.56,23.0)=1.17, p=0.34). (**C**) Paw withdraw latencies on placebo test day from **Extended Figure 1A-B** compared across trials, RM one-way ANOVA with Tukey’s post-hoc (placebo: F(1.99,17.92)=1.85, p=0.19; natural history: F(1.93,17.37)=0.64, p=0.054). (**D**) Schematic of the unpaired placebo conditioning protocol. (**E**) Paw withdrawal latencies on the hot plate across unpaired placebo conditioning and placebo test day (n=10, RM one-way ANOVA with Dunnett’s post-hoc, F(3.25,29.26)=4.33, p=0.011). (**F**) % of morphine effect. (**G**) % responders in the placebo conditioning. (**H**) Distance traveled in the open field for 10-30 minutes after saline, morphine (1^st^ exposure), or placebo (saline) injection, (n=6, RM one-way ANOVA with Tukey’s post-hoc, F(1.83,9.12)=7.377, p=0.014). (**I**) Schematic describing the 30 vs. 10 minute wait period on conditioning and test days, respectively. (**J**) Same as (**A**) but for the scheme described in (**I**) (n=4, RM one-way ANOVA with Dunnett’s post-hoc, F(1.60,4.80)=27.60, p=0.0027). (**K**) Same as (**F**), but for (**I**). (**L**) Same as (**G**), but for (**I**). (M) Hot plate paw withdrawal latencies before and after placebo conditioning with morphine, rc = reconditioned (n=8, RM one-way ANOVA with Dunnett’s post-hoc, F(3.25,22.71)=26.52, p<0.0001). (**N**) Same as (**F**), but for (**M**) (RM one-way ANOVA with Dunnett’s post-hoc, F(2.50,17.52)=3.89, p=0.032). The % morphine analgesia is indicated over each placebo day. (**O**) Same as (**G**), but for (**M**). (**P**) Placebo effect extinction as a function of session number. Data were fit to a monoexponential function.

**Extended Figure 2:**
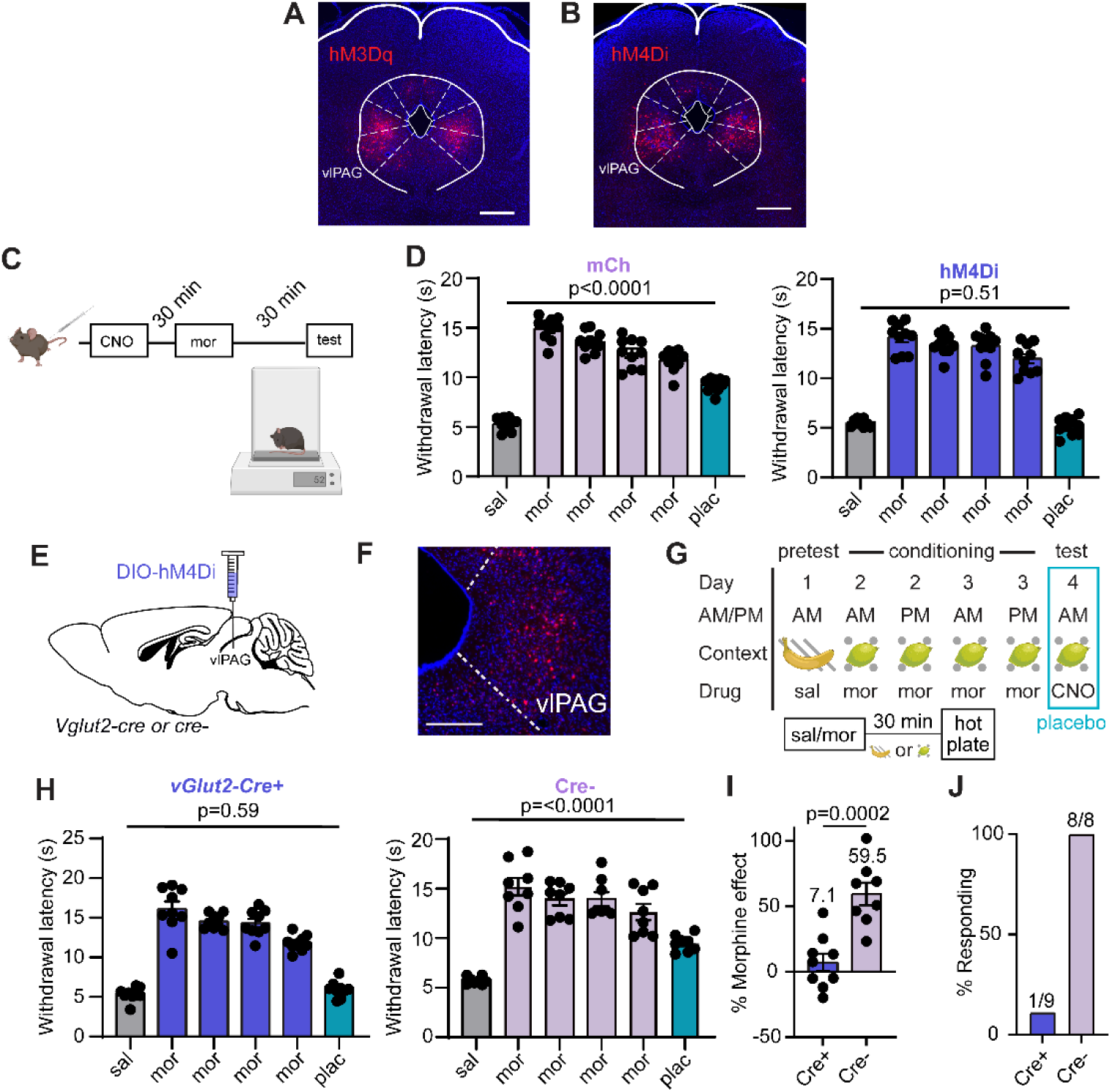
Chemogenetic silencing of vlPAG→RVM and vlPAG*^vGlut2-Cre^* neurons during placebo trials. (**A**) Example of hM3Dq-mCh expression in the vlPAG upon transduction with AAVretro-Cre in RVM, scale bar 0.5 mm. (**B**) Same as **A**, but for hM4Di-mCh. (**C**) Schematic of the timeline for morphine and CNO co-administration to match their onset kinetics for nociception analysis on the hot plate assay in Figure 2F. (**D**) Hot plate paw withdrawal latencies before, during, and after placebo conditioning with morphine upon chemogenetic silencing of vlPAG→RVM neurons on placebo test day for the data shown in Figure 2G-I (mCh: n=10, RM one-way ANOVA with Dunnett’s post-hoc, F(3.31,29.76)=122.8, p<0.0001; hM4Di: n=10, RM one-way ANOVA with Dunnett’s post-hoc, F(3.01,27.07)=133.4, p<0.0001). (**E**) Schematic of viral transduction of vlPAG*^vGlut2-Cre^* neurons with hM4Di. Cre-negative littermates were used as a negative control. (**F**) Example of hM4Di-mCh expression in the vlPAG of a *vGlut2-Cre* mouse, scale bar 0.1 mm. (**G**) Schematic of placebo protocol 2 indicating chemogenetic silencing with CNO administration on placebo test day. (**H**) Hot plate paw withdrawal latencies before, during, and after placebo conditioning with morphine upon chemogenetic silencing of vlPAG*^vGlut2-Cre^* neurons on placebo test day (Cre-: n=8, RM one-way ANOVA with Dunnett’s post-hoc, F(2.79,19.54)=46.28, p<0.0001); Cre+: n=9, RM one-way ANOVA with Dunnett’s post-hoc, F(2.12,16.97)=133.4, p<0.0001)). (**I**) Placebo effect quantified as % of the response to the previous dose of morphine upon chemogenetic silencing of vlPAG*^vGlut2-Cre^* neurons for the data shown in (**H**) (Cre+: n=9, Cre-: n=8, two-tailed t-test). (**J**) Placebo response in (**I**) quantified as the % of subjects exhibiting antinociception.

**Extended Figure 3.**
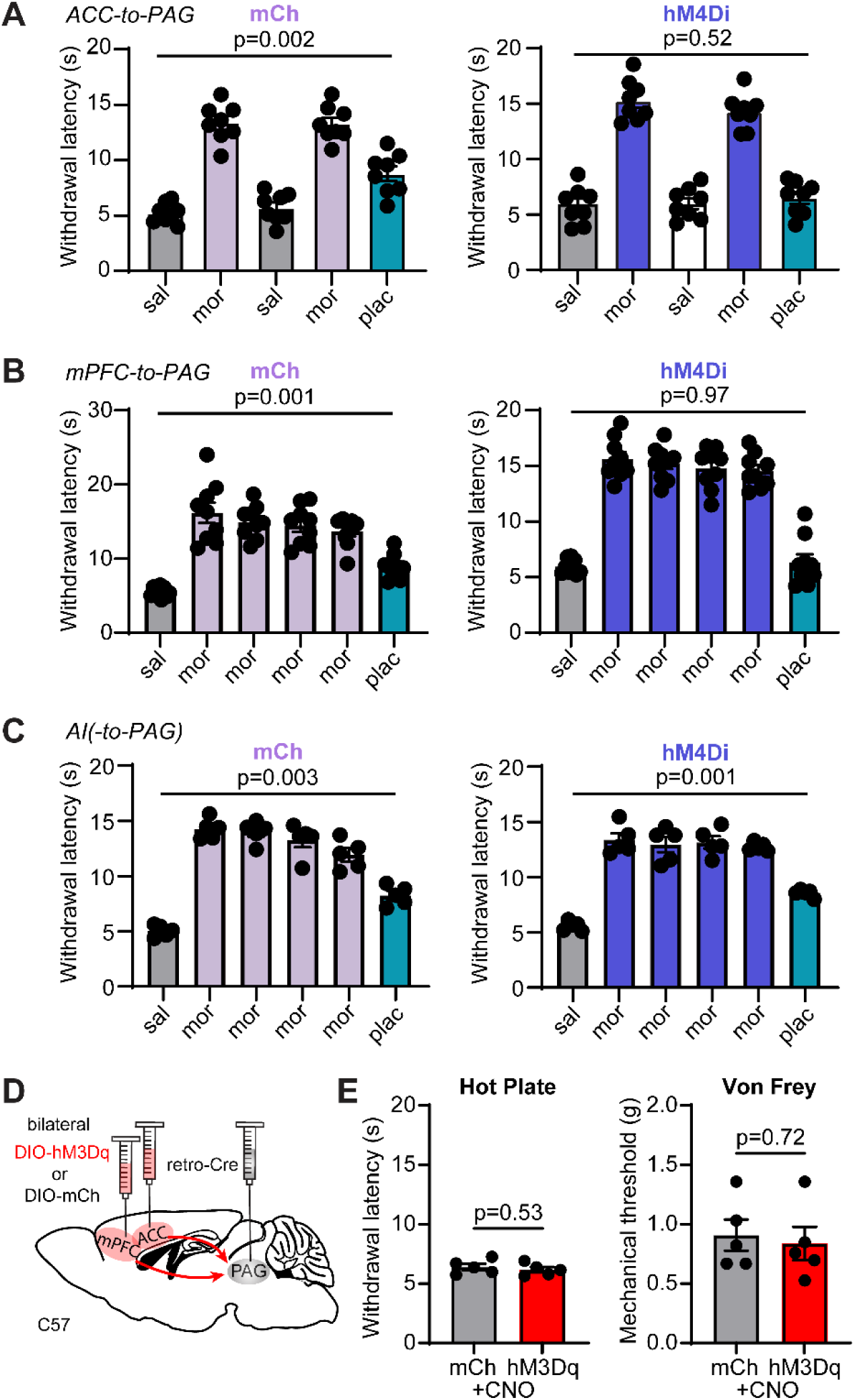
Chemogenetic inactivation of ACC→PAG, mPFC→PAG, and the PAG-projecting region of AI during the placebo test and simultaneous bilateral chemogenetic activation of ACC→PAG and mPFC→PAG neurons. (**A**) Hot plate paw withdrawal latencies before, during, and after placebo conditioning with morphine upon chemogenetic silencing of ACC→PAG neurons on placebo test day (mCh: n=8, RM one-way ANOVA with Dunnett’s post-hoc, F(2.25,15.73)=67.24, p<0.0001); hM4Di: n=8, RM one-way ANOVA with Dunnett’s post-hoc, F(2.42,16.92)=93.54, p<0.0001). (**B**) Hot plate paw withdrawal latencies before, during, and after placebo conditioning with morphine upon chemogenetic silencing of mPFC→PAG neurons on placebo test day (mCh: n=9, RM one-way ANOVA with Dunnett’s post-hoc, F(2.08,16.63)=63.68, p<0.0001); hM4Di: n=9, RM one-way ANOVA with Dunnett’s post-hoc, F(2.65,21.23)=113.8, p<0.0001). (**C**) Hot plate paw withdrawal latencies before, during, and after placebo conditioning with morphine upon chemogenetic silencing of the PAG-projecting region of AI on placebo test day (mCh: n=5, RM one-way ANOVA with Dunnett’s post-hoc, F(3.23,12.90)=126.9, p<0.0001); hM4Di: n=5, RM one-way ANOVA with Dunnett’s post-hoc, F(2.09,8.37)=59.75, p<0.0001). (**D**) Schematic of the retrograde viral injection approach for chemogenetic activation of ACC→PAG and mPFC→PAG neurons. (**E**) Paw withdrawal latencies on the hot plate (left) and mechanical thresholds to stimulation with von Frey fibers (right) upon chemogenetic activation of both ACC→PAG and mPFC→PAG neurons with hM3Dq vs. mCherry control (mCh: n=5, hM3Dq: n=5, unpaired t-tests).

**Extended Figure 4:**
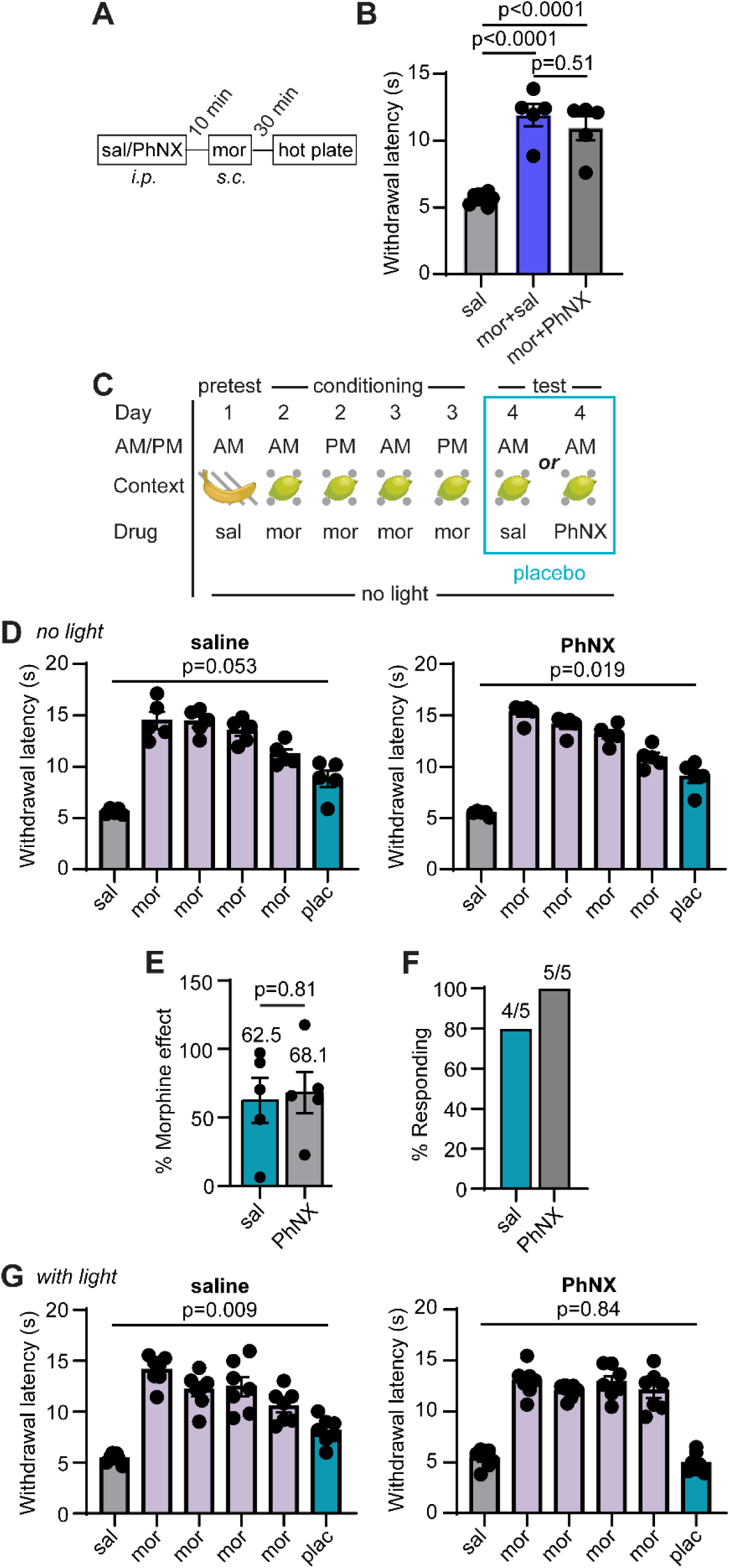
PhNX is inactive in the absence of light. (**A**) Schematic of PhNX (30 mg/kg *i.p.*) and morphine (5 mg/kg *s.c.*) administration to assess potential antagonism of opioid signaling by PhNX *in vivo*. (**B**) Paw withdrawal latencies on the hot plate in response to saline, morphine and saline, or morphine and PhNX (saline: n=10, mor+sal: n=5, mor+PhNX: n=5, One-way ANOVA with Tukey’s post-hoc, F(2,17)=44.81, p<0.0001). (**C**) Schematic of Placebo protocol 2 with administration of either saline or PhNX (30 mg/kg *i.p.*) on test day in the absence of photoactivation. (**D**) Hot plate paw withdrawal latencies before, during, and after placebo conditioning with morphine in the absence and presence of PhNX without light on placebo test day (sal: n=5, RM one-way ANOVA with Dunnett’s post-hoc, F(1.83, 7.30)=30.18, p=0.0003); PhNX: n=5, RM one-way ANOVA with Dunnett’s post-hoc, F(2.16,8.64)=76.96, p<0.0001). (**E**) Placebo effect quantified as % of the response to the previous dose of morphine (saline: n=5, PhNX: n=5, unpaired t-test). (**F**) Placebo response quantified as the % of subjects exhibiting antinociception. (**G**) Hot plate paw withdrawal latencies before, during, and after placebo conditioning with morphine in the absence and presence of PhNX and light on placebo test day (sal: n=7, RM one-way ANOVA with Dunnett’s post-hoc, F(2.39,14.34)=26.91, p<0.0001); PhNX: n=7, RM one-way ANOVA with Dunnett’s post-hoc, F(2.91,17.47)=69.55, p<0.0001).

